# Single molecule sequencing of THCA synthase reveals copy number variation in modern drug-type *Cannabis sativa* L

**DOI:** 10.1101/028654

**Authors:** Kevin J. McKernan, Yvonne Helbert, Vasisht Tadigotla, Stephen McLaughlin, Jessica Spangler, Lei Zhang, Douglas Smith

## Abstract

- Cannabinoid expression is an important genetically determined feature of *cannabis* that presents clinical and legal implications for patients seeking cannabinoid specific therapies like Cannabidiol (CBD).
- Cannabinoid, terpenoid, and flavonoid marker assisted selection can accelerate breeding efforts by offering genetic tools to select for desired traits at an early stage in growth. To this end, multiple models for chemotype inheritance have been described suggesting a complex picture for chemical phenotype determination.
- Here we explore the potential role of copy number variation of THCA Synthase using phased single molecule sequencing and demonstrate that copy number and sequence variation of this gene is common and suggests a more nuanced view of chemotype prediction.

## Introduction

The genetics of cannabis chemotype has been extensively studied due to the highly selected THCA and CBDA phenotypes. A co-dominant model of inheritance of THCA and CBDA synthase (THCAS, CBDAS) (de Meijer *et al*., 2003) has been demonstrated describing a Bt:Bd allele in linkage to the synthase genes (Onofri *et al*., 2015; Weiblen *et al*., 2015). In addition to this Bt:Bd allele, single nucleotide polymorphisms in the FAD binding domain of THCAS have been described that impair the function of the enzyme (Sirikantaramas *et al*., 2004). Cascini *et al*. demonstrated varying copy number of a highly conserved region in THCAS but did not find this assay to be correlative to THCAS expression (Cascini *et al*., 2012; Cascini *et al*., 2013). Complimenting de Meijer and Sirikantaramas work, van Bakel *et al*. reported a gene of unknown function (AAE3) that was differentially replicated in drug versus fiber type cannabis (van Bakel *et al*., 2011).

Whole genome sequencing using 454 and Illumina sequence data demonstrated excessive polymorphic coverage over the single exon THCAS gene(McKernan, https://archive.org/details/SequencingTheCannabisGenome)) but allelic phasing of these important genes has not been possible using short read sequencing and conventional genome assembly technologies. Using bacterial cloning, Onofri *et al*. demonstrated a polyploid status for THCA synthase (THCAS) confirming the putative copy number variation observed with whole genome sequencing. Nevertheless, this polyploid status appeared to be a rare event occurring in a single United States bred cultivar and was not correlated with increased THCA expression. Weiblen *et al (Weiblen et al., 2015).* and Cascini *et al. (Cascini et al., 2012; Cascini et al., 2013)* further supported polyploidy with quantitative PCR and bacterially cloned THCAS DNA but deep sampling of alleles may have been limited by bacterial cloning methods. Here, we apply over 10,000 long-read single molecule sequences per sample in combination with Illumina sequencing methods, to phase the multiple copies of THCA synthase in over a dozen modern medicinal cultivars (Eid *et al*., 2009). Allelic phasing of this 1.6kb gene adds further support to additional mechanism of polyploidy in chemical inheritance.

Multiple primer pairs were designed to amplify THCAS, and the amplicons were sequenced using Pacific Bioscience’s circular consensus sequencing (CCS) method and a size-selected Illumina paired-250bp read system (Table1). Given the 40kb reads of the Pacific Biosciences (PacBio) platform, it is ideally suited for phasing polymorphic genes that are longer than the read lengths accessible by Sanger sequencing. Bacterial cloning and primer walking has been used traditionally to address this, but the methods can be cumbersome, expensive and are susceptible to cloning bias (Metzker, 2010). Single molecule sequencing enables one to deeply sequence a PCR product without bacterial cloning to eliminate this bias. The higher raw error rate of single molecule sequencing is overcome by ligating the target amplicon into a circle and performing 20X rolling circle sequencing (Chaisson *et al*., 2015). This CCS procedure significantly reduces the stochastic error associated with single molecule detection while providing long phased haplotypes of each copy of the amplified gene. These phased haplotypes were then verified by paired-end Illumina sequencing to confirm pure phase with homozygous alignments. Cultivars that vary in CBDA and THCA content were sequenced to assess the diverse haplotypes present in *cannabis* and to resolve putative pseudogenes from intact open reading frames.

**Table1.**
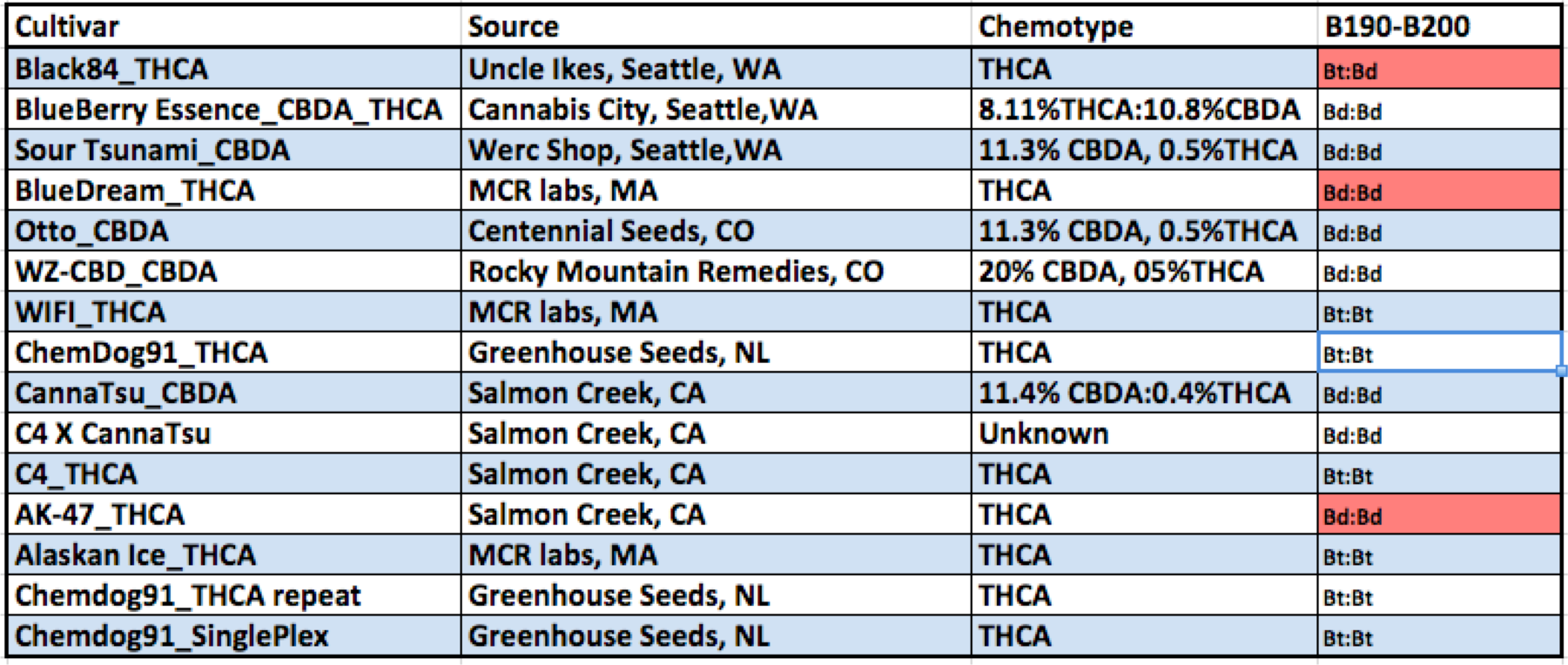
THCAS was amplified with two primer sets (Onofri and MGC-2130) for each Cultivar and single molecule sequenced with Pacific BioSciences SMRT cell circular consensus sequencing. B190, B200 alleles were measured according to Mandolino *et al*. Red highlighted samples represent B190-B200 Genotype-Chemotype discordant samples.

| Cultivar | Source | Chemotype | B190-B200 |
| --- | --- | --- | --- |
| Black84_THCA | Uncle Ikes, Seattle, WA | THCA | Bt:Bd |
| BlueBerry Essence_CBDA_THCA | Cannabis City, Seattle, WA | 8.11%THCA:10.8%CBDA | Bd:Bd |
| Sour Tsunami_CBDA | Werc Shop, Seattle, WA | 11.3% CBDA, 0.5%THCA | Bd:Bd |
| BlueDream_THCA | MCR labs, MA | THCA | Bd:Bd |
| Otto_CBDA | Centennial Seeds, CO | 11.3% CBDA, 0.5%THCA | Bd:Bd |
| WZ-CBD_CBDA | Rocky Mountain Remedies, CO | 20% CBDA, 05%THCA | Bd:Bd |
| WIFI_THCA | MCR labs, MA | THCA | Bt:Bt |
| ChemDog91_THCA | Greenhouse Seeds, NL | THCA | Bt:Bt |
| CannaTsu_CBDA | Salmon Creek, CA | 11.4% CBDA:0.4%THCA | Bd:Bd |
| C4 X CannaTsu | Salmon Creek, CA | Unknown | Bd:Bd |
| C4_THCA | Salmon Creek, CA | THCA | Bt:Bt |
| AK-47_THCA | Salmon Creek, CA | THCA | Bd:Bd |
| Alaskan Ice_THCA | MCR labs, MA | THCA | Bt:Bt |
| Chemdog91_THCA repeat | Greenhouse Seeds, NL | THCA | Bt:Bt |
| Chemdog91_SinglePlex | Greenhouse Seeds, NL | THCA | Bt:Bt |

## Results

We demonstrate that multiple copies of THCAS with distinct sequences are present in many common dispensary and coffee house cultivars in circulation in 2015 (Figure 1). The ascertainment of THCAS copy number is dependent on primer selection suggesting local divergence in the 5’ and 3’ untranslated regions (UTRs).

**Figure 1.**
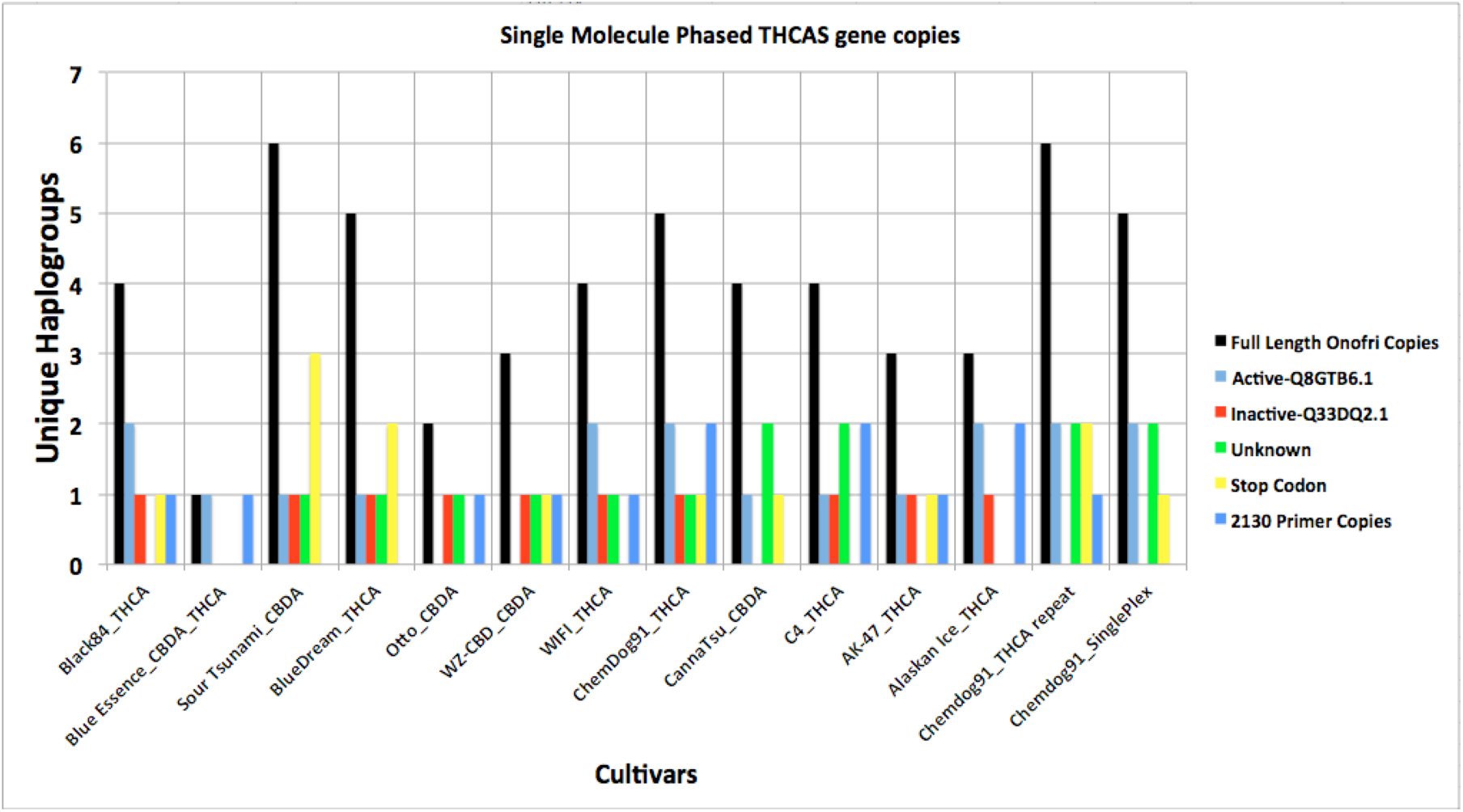
Unique full length THCA Synthase sequences for each cultivar demonstrate diploidy is a rare event in Drug Type cultivars selected for high cannabinoid content. All THCA positive strains have an active form of THCAS with the exception of AK-47 that has an A250D variant of the active form Q8GTB6.1. A250D variants are also present in Black84 as the only possible functional THCAS and as the 2^nd^ THCAS haplogroup in Chemdog91. While it is also present in a CannaTsu high CBDA cultivar, this cultivar does not amplify with the MGC-2130 primers suggesting an MGC-2130 A250D haplogroup is in fact an active THCAS when full length.

Utilizing two sets of primer pairs (within and external to THCAS), we amplified polyploid copies of THCAS and sequenced them using the Pacific Biosciences CCS method. To mitigate PCR error we required a minimum of 10 independent, but identical sequences from different zero mode waveguides (ZMW) to establish a haplogroup. For each CCS, the molecule was read on the forward and reverse strands 20-30 times to form a circular consensus that was over 99.95% accurate (Table 2).

**Table 2.**
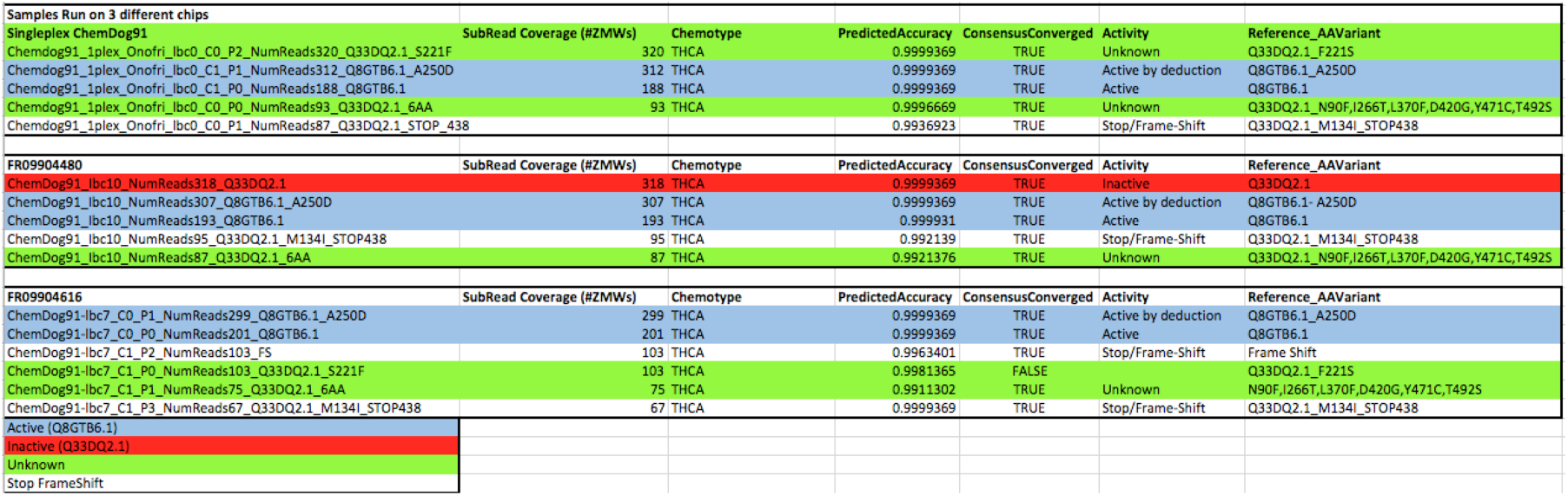
Triplicate Sample analysis. ChemDog91 was amplified 3 distinct times and sequenced on 3 different SMRT cells to assess reproducibility and sampling. Active forms consistently amplify and sequence across all amplification products. A few private inactive or unknown activity haplogroups amplify inconsistently (Q33DQ2.1-Red and Q33DQ2.1_F221S-Green).

Once PacBio long reads were error corrected using CCS, Illumina Nextera libraries were generated from size-selected 850 bp fragments derived from the PCR products, and were sequenced with 250 base paired-end reads. The resulting read pairs could be accurately mapped to each PacBio haplogroup to produce homozygous alignments for each allele (Figure 2.). These homozygous alignments were helpful in ascertaining whether all haplogroups were accurately represented in PacBio data and if there were any remaining variants ambiguously mapped. Unphased polymorphic data is difficult to translate into amino acid sequence and obscures accurate assessment of functional alleles.

**Figure 2.**
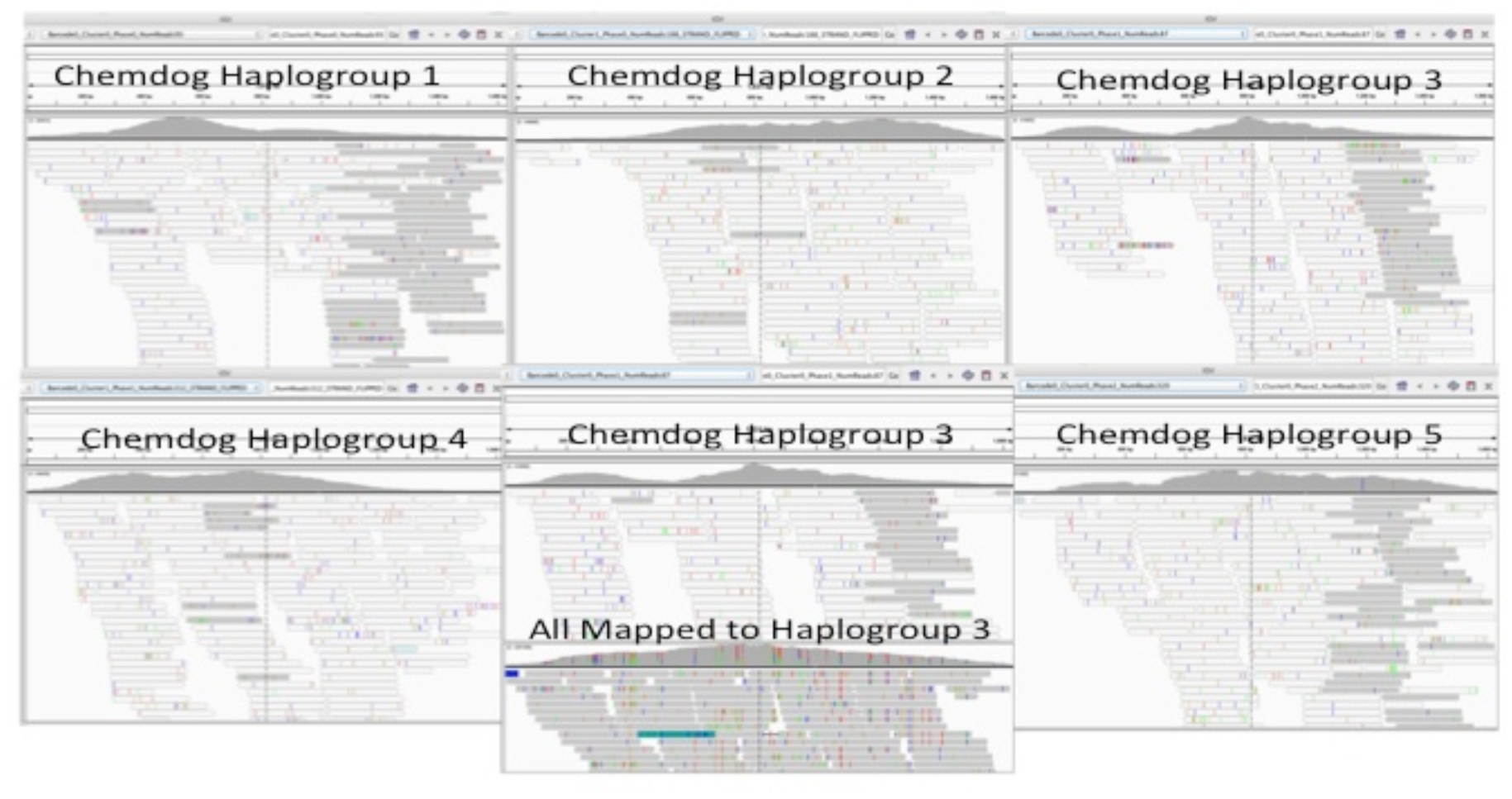
Mapping 2x250 Illumina reads to all of the Phased Pacific Bioscience generated haplogroups compared to mapping them to a single THCAS haplogroup sequence as you might find in CanSat3. 5 haplogroups of Chemdog91 shown in IGV with Illumina reads mapped to a refernce containing all 5 references. The center bottom view “All Mapped to Haplogroup3” represents all of the reads mapped to just Haplogroup 3 demonstrating collapsed polymorphic reads when mapped to the unphased reference sequence of a single haplogroup. Most reads in IGV are labeled with poor mapping scores due to homology between the haplogroups. IGV also filters out alleles under 20% frequency inducing a more detailed dissection of this across more samples in Figure 5.

The THCAS amplicons were initially amplified using the primers described by Onofri *et al. (Onofri et al., 2015)* This primer set spans the start and stop codons of the THCAS gene and thus is incapable of providing sequence information for the first and last 25bp of the gene. Moving the primers out into the flanking non-coding regions (MGC-2130 external primer set) delivered fewer haplogroups for all cultivars tested, presumably due to the divergence of these regions in the different gene copies (Figure 3).

**Figure 3.**
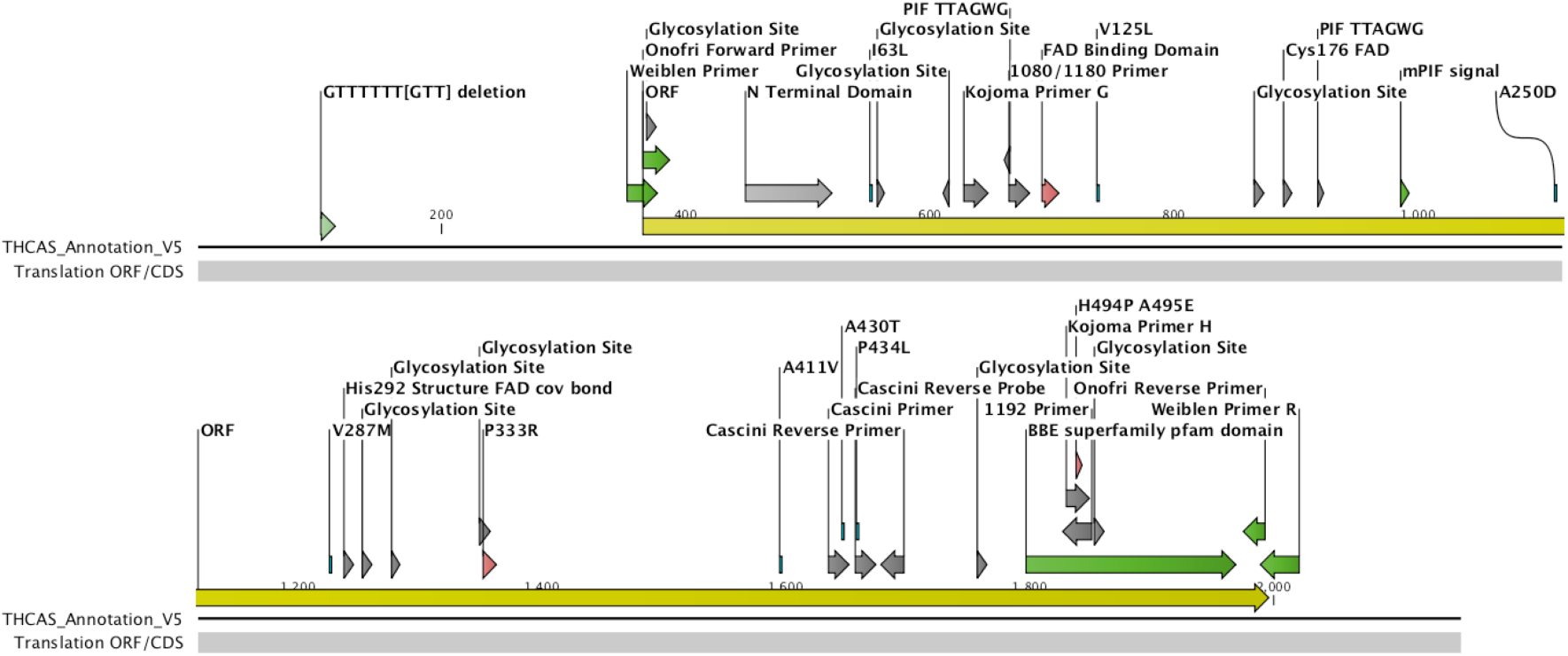
THCA Synthase Annotation. MGC-2130 Primers bracket the displayed region of interest. Onofri and Weiblen primers are in green. THCAS ORF is in yellow. Various nonsynonmous SNPs (I63L, V125L, A250D, V287M, M289T, P333R, A411V) found in Active THCAS haplogroups are annotated near respective functional groups like Glycosylation sites, FAD binding domains and mPIF signals. Cascini primers and BBE Pfam domains are labeled accordingly.

To ascertain the reproducibility of this approach, Chemdog91 was amplified 3 different times and run on 3 different SMRT cells. One sample was run as a single-plex sample on a SMRT cell to afford high enough sampling to ensure no alleles were missed. While each sample consistently sequenced the active haplogroups to high coverage, a few inactive or unknown haplogroups emerged on only 2 of the 3 SMRT cells. Since one of the intermittent haplogroups has a unique variant (S221F genotype) private to Chemdog91, we can rule out contamination from other gDNA being the source of these intermittent haplogroups. We cannot rule out the possibility of other diverged copies of inactive THCAS stochastically amplifying given the PCR reactions were set up independently. Gradient PCR at lower annealing temperatures with the Onofri primers may unveil additional diverged pseudogenes. A few intermittent haplogroups had depressed consensus accuracy and are detectable with appropriate quality filtering tools. As a result of these conditions we do not believe the amplicon subread coverage numbers derived from the Onofri primer sets can be taken as a direct proxy for absolute copy number in the genome. It is possible that some of the high coverage, active THCAS haplogroups may be more than diploid in copy number yet exist as non-diverged sequence in the genome. With the MGC-2130 data being multiplexed, any inter-strain coverage or inferred copy number differences are more likely the result of normalization of barcoded samples prior to multiplexing. Only intra-strain copy number can be assessed with these data.

We segregated the haplogroups based on two genbank accession sequences that have been confirmed as encoding active (Q8GTB6.1 also identical to genbank accession number E33090) and inactive (Q33DQ2.1) THCAS (Sirikantaramas *et al*., 2004; Kojoma *et al*., 2006). While these two haplogroups differ by 37 amino acids from each other, several other haplogroups diverged by up to 2 and 6 amino acids from these two reference sequences (Figure 4, Table 3). Other haplogroups were identified that contained termination or frame-shifting variants. These had higher similarity to Q33DQ2.1 than Q8GTB6.1, suggesting an inactive heritage. Phylogenetic trees were constructed that clustered the divergent haplogroups around Active, Inactive, and frame-shifted clusters (Figure 4).

**Figure 4.**
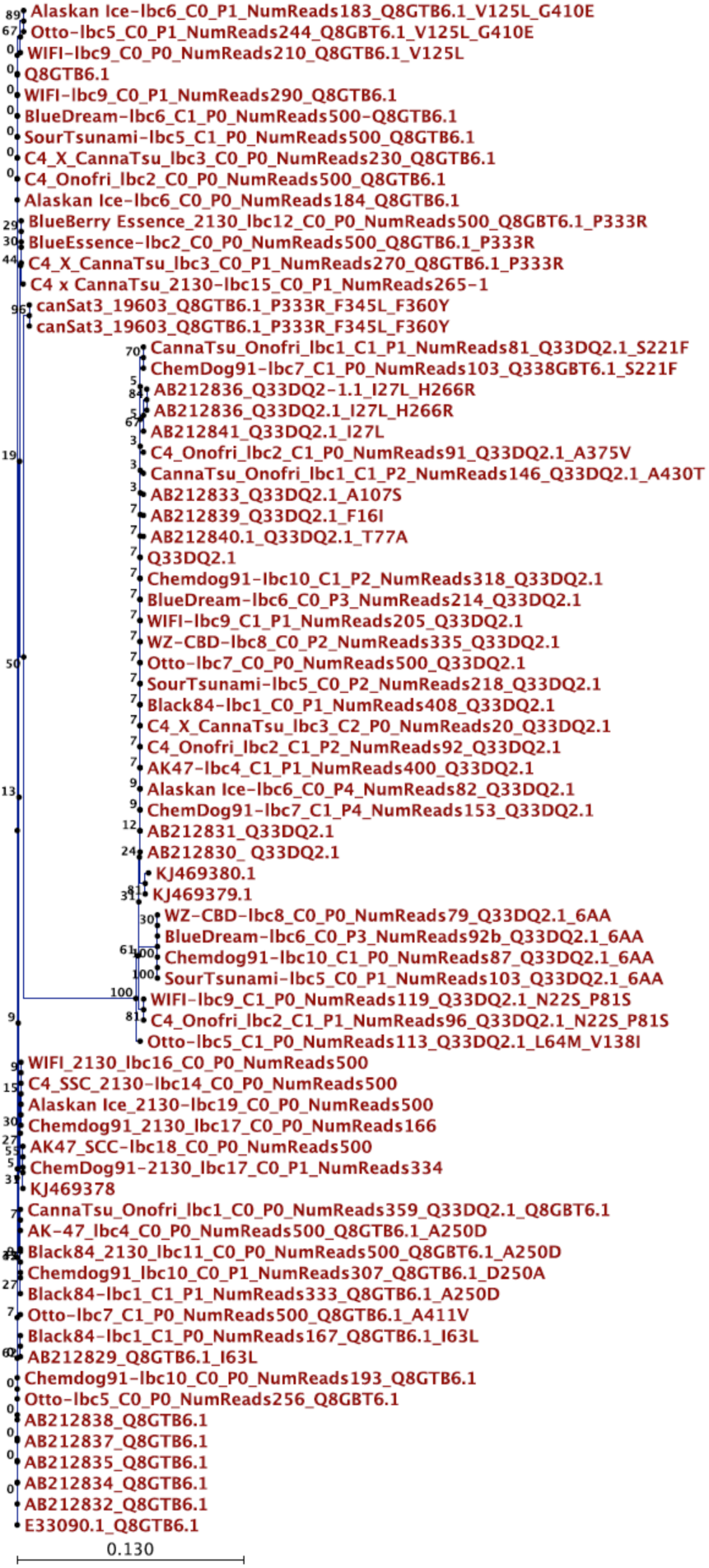
Phylogenetic tree of amino acid sequences of THCAS using E33090 as the root. Long pseudogenes were omitted from the tree since they had excessive divergence. The NumReads reflects the number of subreads that support a haplogroup after circular consensus has been achieved. Most haplogroups cluster around Active and Inactive like accessions (Q8GTB6.1 & Q33DQ2.1 accessions). Only Q8GTB6.1 cluster amplify with MGC-2130 primers. Other GenBank Accessions are label as AB#.

**Figure 5.**
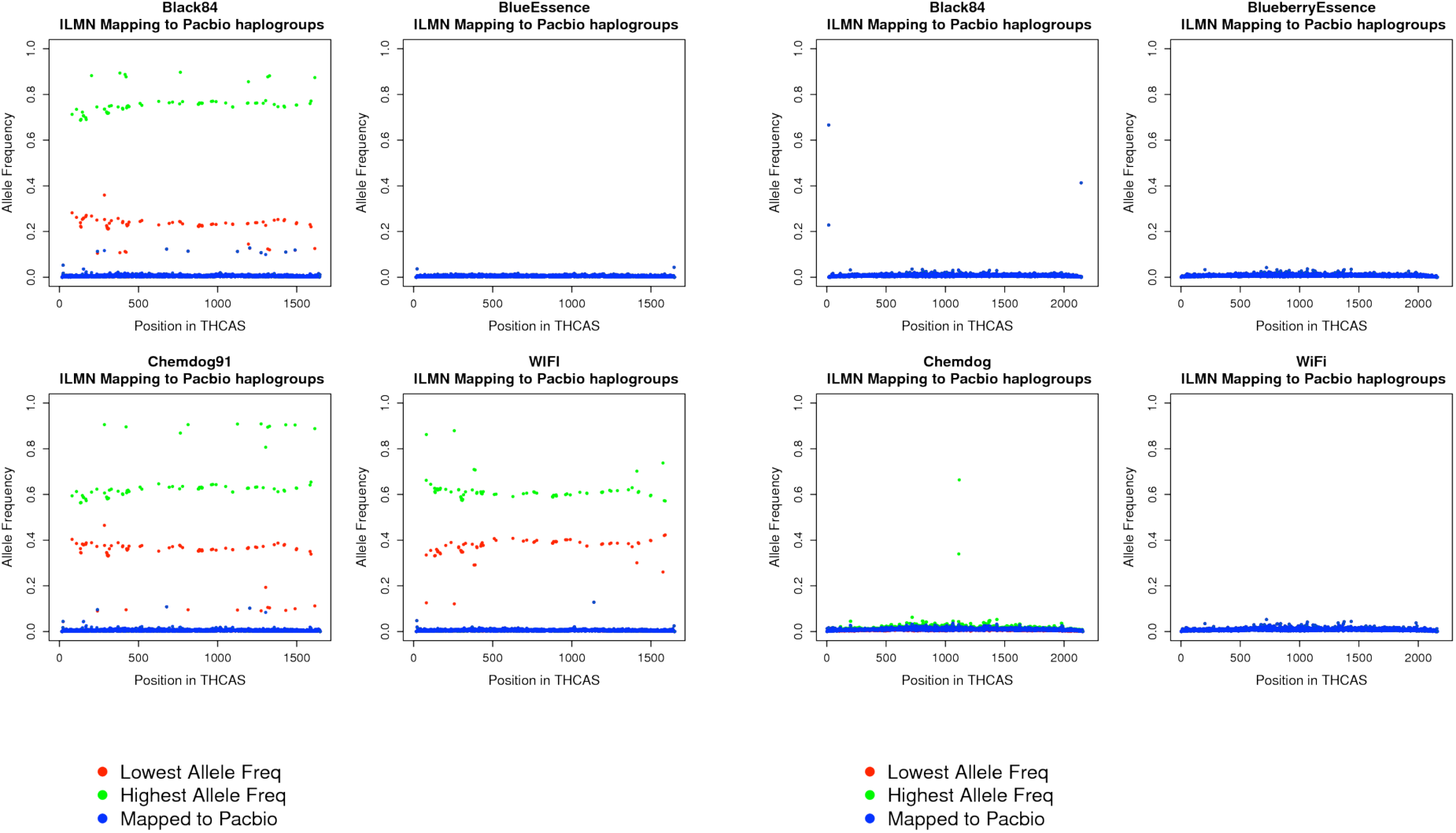
Over 100,000X coverage with Illumina 2x250bp reads derived from Onofri and MGC-2130 primer based amplification of THCAS. The Illumina data is mapped to the Pacbio FASTA files derived from Pacbio LAA analysis. The strain sequence is respectively mapped- for example- *Black84* Illumina Data is mapped to the *Black84* Pacbio FASTA. As an example, if there are 5 haplogroups, the data is mapped 6 times; once to each separate haplogroup and once to all 5 together in one FASTA reference. An allele count is performed on each position of THCAS. A clustal alignment is performed between all the haplogroups so we are able to compare the same position across the multiple reference/bam files. The minimum and maximum non-reference ratio at each position (~1600) is calculated across all the haplogroups (i.e. 5). This is the green and red dots shown. The non-reference ratio for all haplogroups mapped together is also calculated and shown in blue. Left two columns of the chart is Onofri amplification allelic coverage charts. Right 2 columns are MGC-2130 amplicon allelic coverage charts. On the X-axis is THCAS bases 1-1654bp or 1–2130bp respectively. On the Y-axis is Heterozygosity where homozygous genotypes are either 1 or 0 and heterozygous genotypes would display at 0.5. Samples Blueberry essence has been fully sampled with the PacBio depth on both primer sets (all blue homozygous alignments). Samples with red and green dots at equal allele frequency across the amplicon represent PacBio data that may have a remaining haplogroups detected at 100,000 X Illumina sequence but not yet sampled at the PacBio read depth. All 2130-MGC amplicons look fully sampled with phased active alleles.

**Table 3.**
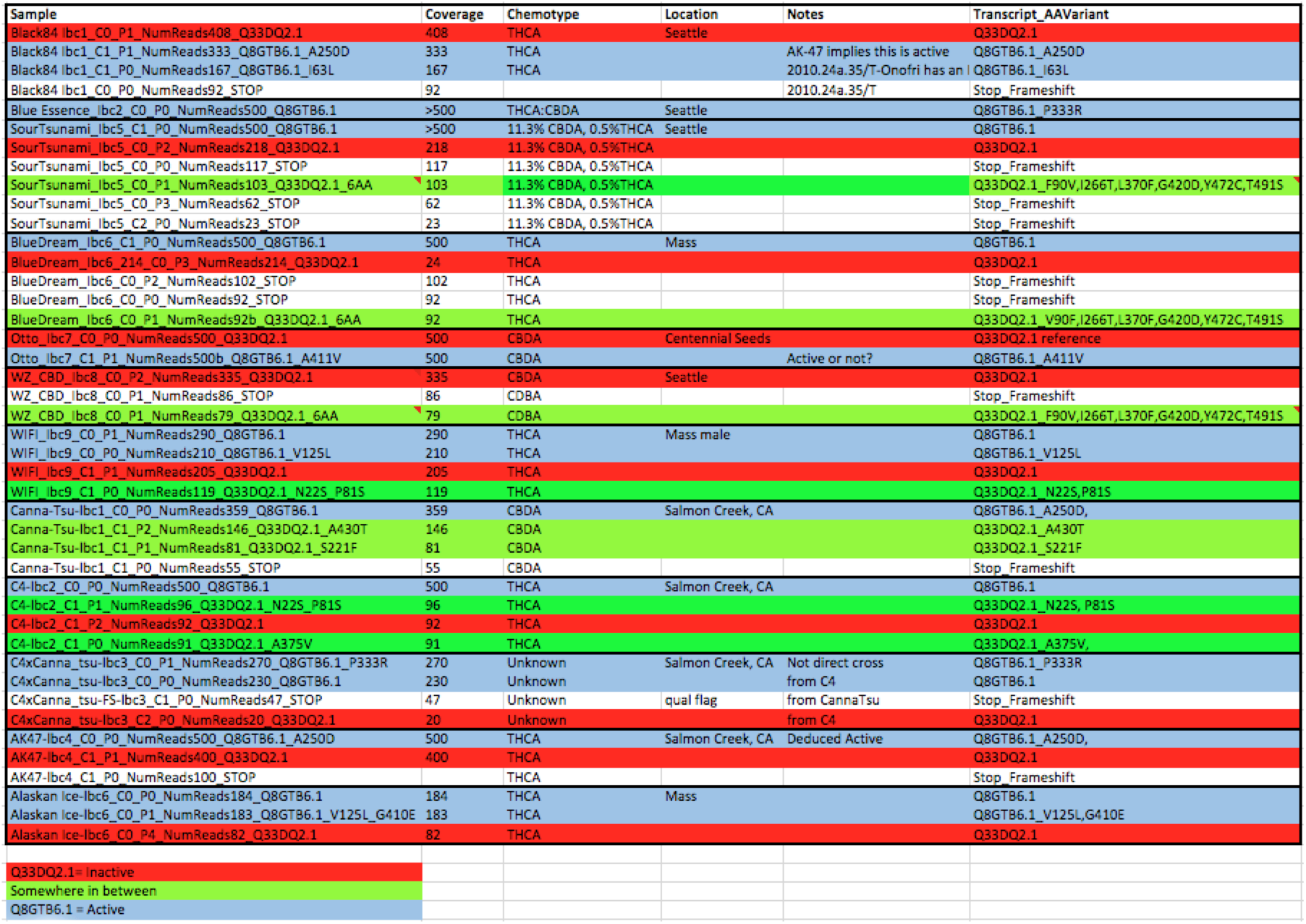
List of all haplogroups discovered from each cultivar other than Chemdog91 (Table 2). Red highlighting are Inactive haplogroups. Green highlightings are unknown haplogroups and blue highlightings are active haplogroups.

We surveyed CBDA and THCA dominant cultivars seen in Table 1, 2 and 3. Cultivars measured to have CBDA-rich chemotypes like WZ_CBD contained 3 haplogroups, one of which was clearly a pseudogene (Figure 1). The other two haplogroups were either identical to, or had 6 amino acid mismatches with the “Inactive THCAS” in genbank (Q33DQ2.1). Since this cultivar has low but measured THCA (0.6% THCA: 20%CBDA), it could be assumed that the 6 amino acid divergence from the inactive form represents a very weakly active THCAS gene. The cultivar was positive for the Bd:Bd genotype described by de Miejer, and if this allele governs the binary expression of CBDAS over THCAS then the expression level of THCAS is uncertain and the impact of the 6 amino acid variants on activity is impossible to infer. Some level of chemotype expression exists and suggests a leaky Bd:Bd expression is responsible for the 0.6% THCA. Of note, this haplogroup failed to amplify with the MGC-2130 external primer set (Table 4).

**Table 4.**
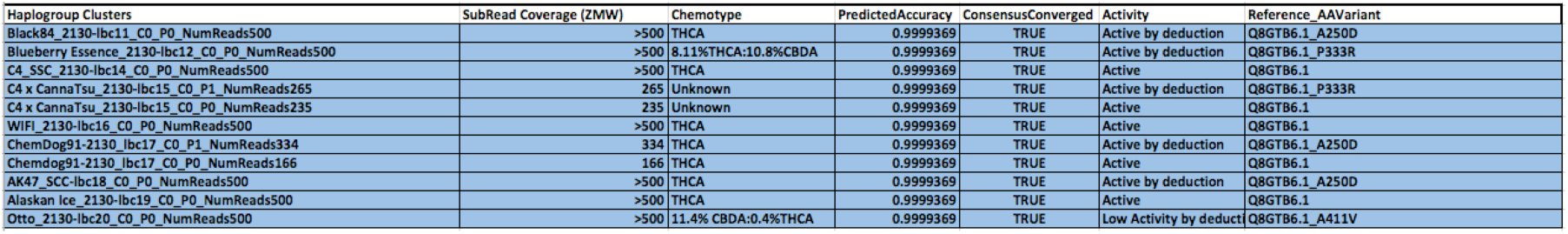
MGC-2130 primers only amplify active alleles. Active transcript Q8GTB6.1 is the most common haplogroup sequenced. A250D is the second most common haplogroup and is the only haplogroup present in a high THCA expressing cultivar AK-47 and is thus labeled as “Active by deduction”. P333R is also present in Blueberry Essence with 8.11% THCA suggesting Active by deduction Classification. A411V may be low activity or in a cultivar with Bd:Bd genotype.

The 6 amino acid diverged THCAS haplogroup is also found in other THCA dominant cultivars like Chemdog91, however Chemdog91 is a high THCA expressing cultivar and has 5 THCAS copies in total (one with a stop codon, one that is inactive and three that are putatively active). One haplogroup in Chemdog91 is identical in amino acid sequence to the active THCAS sequence (Q8GTB6.1) and the other has a common A250D variant of the active form. Although we did not collect RNA from these samples, Onofri, *et al*. also describe some of these alternative alleles and demonstrates RNA expression (Onofri *et al*., 2015).

Adding to the observations of Onofri, *et al*., we were able to find THCA producing strains that exclusively held the ambiguous genotypes described in their paper as (1/4, 2/1, 2/2, 2/3). The 1/4 genotype described by Onofri, *et al*. is an A250D variant of the active form. The Cultivar “AK-47” presented with 3 haplogroups of which one had a stop codon, one had an inactive allele (Q33D12.1) and one presented with a single active THCAS haplotype with an A250D variant. We can deduce from this that the A250D allele is the only haplogroup capable of synthesis of the high levels of THCA found in AK-47.

Interestingly, another high THCA strain (Black84) shared this A250D variant but also had the I63L variant described and deemed active by Onofri et al. (Table 3). Only the A250D variant amplified with the MGC-2130 primer set bringing some question to the activity of the I63L variant in our cultivars. Chemdog91, in addition to its Q8GTB6.1 active allele, also has an A250D haplogroup. Both of these haplogroups amplify with the MGC-2130 primer set in Chemdog91 and only the A250D allele amplifies in Black84 suggesting the I63L variant may be inactive (Table 4). Further work is required to characterize the I63L variants functionality.

The other THCA:CBDA hybrid cultivar “Blueberry Essence” has a single haplogroup that amplified with both primers, suggesting the sample had identical maternal and paternal alleles for THCAS sequence. This sequence has a single amino acid variant from the active form (P333R). Considering the structure of the gene, this proline change may alter the activity of the THCAS and explain the 8.11% THCA, 10.8%CBDA expressed in Blueberry Essence (Shoyama *et al*., 2012). Figure 6 demonstrates a Hybrid Bd:Bd genotype. The difference in Bd:Bd vs Bt:Bd genotypes relies on measuring the magnitude of the 3^rd^ peak in the electropherogram but in our hands the assay did not produce a clear presence-absence result allowing unambiguous differentiation of these two alleles (Figure 6.). Interestingly, the B1080/B1192 primers described in Pacifico *et al*. were confirmed to be the B1180/B1192 genotyping primers utilized in Table 2 of Onofri *et al*. (personal communication G. Mandolino). This forward primer targets a region conserved between active and inactive THCAS while the mid portion of the reverse primer rests on a H494P and A495E variant between active and inactive alleles. Although the 3 prime end of the primer is 12 bases upstream of these primed variants, it is possible but unlikely that this primer set differentiates between active and inactive transcripts of THCAS.

**Figure 6.**
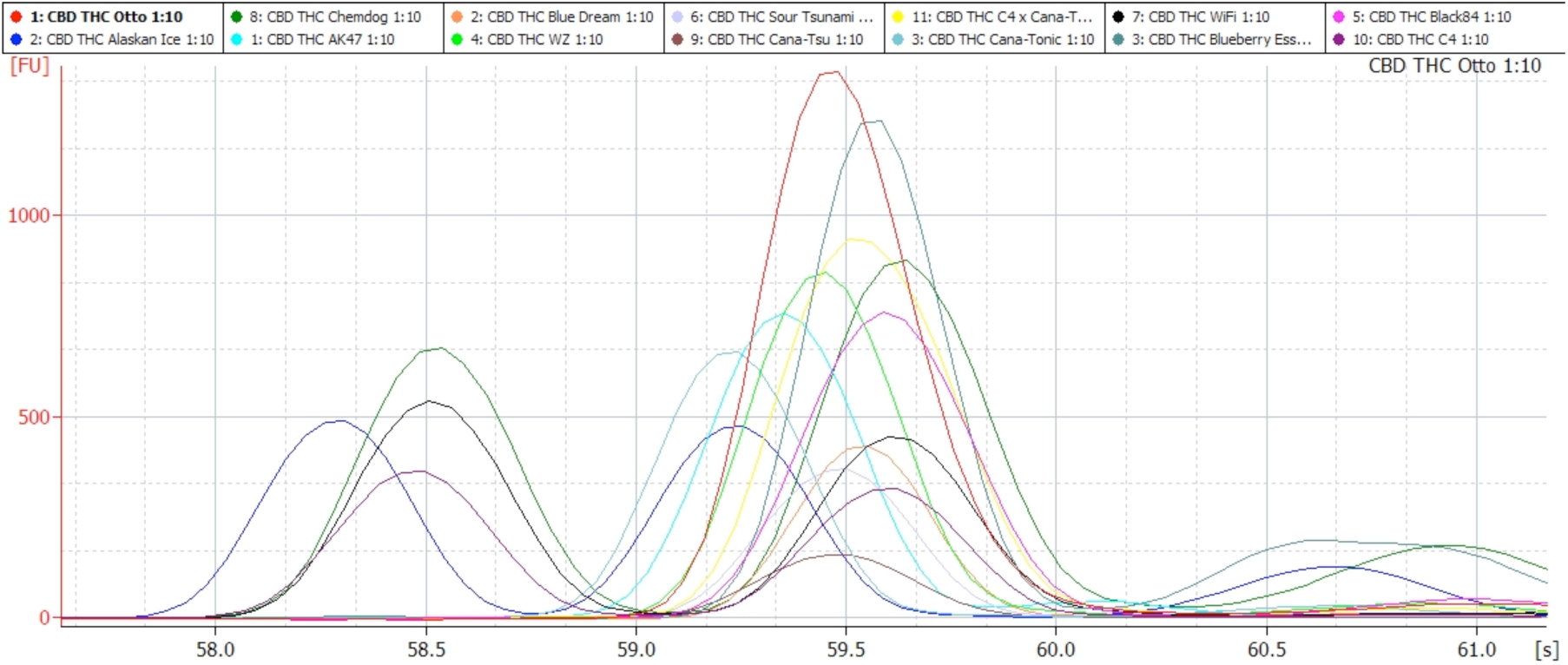
Bt:Bd allele amplified and run on an Agilent High Sensitivity chip. High THCA cultivars (WIFI, Chemdog, C4, Alaskan Ice) replicate a small ~190bp band known as Bt:Bt. High CBDA alleles generate a major 200 bp and a minor 215bp band known as Bd:Bd. Hybrid Bt:Bd alleles generate similar bands as Bd:Bd but with more 215bp product. This later peak is a bit subjective to measure.

Otto and Sour Tsunami are both CBDA dominant chemotypes but demonstrate 2 and 6 haplogroups respectively. Three of the haplogroups in Sour Tsunami have stop codons. The remaining three haplogroups are exact active and inactive haplogroups with an additional 6 amino acid variant of the inactive haplogroup with unknown activity. Otto has 2 haplogroups, one inactive copy that only amplifies with Onofri primers and another that amplifies with both primer sets and represents a single active Q8GTB6.1 haplogroup with an A411V variant.

## Illumina Sequencing of the Bt:Bd alleles

We amplified the Bt:Bd alleles from all of the samples in Table 1 (Figure 6). While 4 of the THCA positive samples demonstrated the 190bp band, 3 did not. A select few of these products were Illumina sequenced to gain a better understanding of the loci the B190:200 primers amplify. Assembling these amplicons produced several contigs per cultivar where the inserts can be mapped to a tandem repeat in CanSat3 Scaffold 19079 (Figure 7.). Only 9-10bp of the 3’ end of the B190/B200 primers match the reference at appropriate distances implying a Tm sensitive assay that is likely highly sensitive to salts in various DNA isolations. Of note, the referenced methods in Mandolino et al. recommend 38C annealing temperatures (Mandolino, 1999). We confirmed these alignments with cross comparisons to other public whole genome assemblies with of Chemdog91 and LA Confidential(McKernan, https://aws.amazon.com/datasets/the-cannabis-sativa-genome/). Contig_60162 of LA Confidential has a 100% 159bp alignment of WZ-CBD B190/B200 allele while Chemdog91s contig_96784 has a 148/159bp alignment with many other contigs showing similar homology. Various polymorphisms also exist in the alignments. While this appears to be a repeat, it is also rich in secondary structure notorious for error prone sequencing (Nakamura *et al*., 2011).

**Figure 7.**
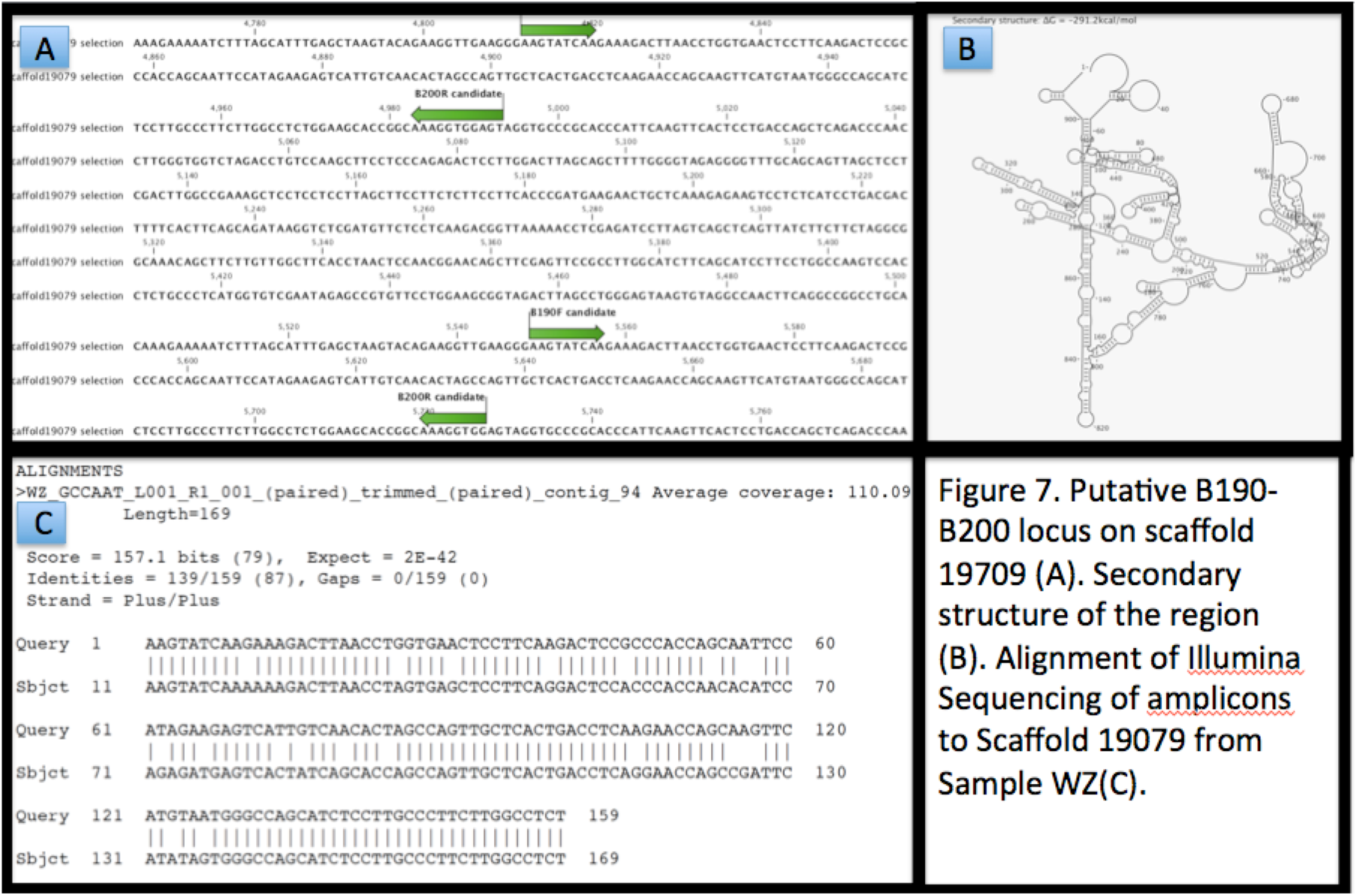
Assembly of Illumina reads from Bt:Bd allele in Cultivar WZ with homology to the B190/B200 partial primer sequences highlighted in green arrows.

While these are useful markers for tracking chemotype, its unclear by their sequence alone what function they play. Even though THCAS rests on CanSat3 scaffold 19603 and shares little homology to CanSat3 scaffold 19079, one possibility is that the regions are simply in linkage to active and inactive alleles of CBDAS and THCAS and closer markers to the gene may be more predictive in a polymorphic population.

## Discussion

Even though absolute copy number does not correlate with CBDA dominant or THCA dominant chemotypes, a more nuanced view of the amino acid alterations in the haplogroups reveals an interesting pattern. All CBDA strains contain at least one haplogroup with similarity to the inactive Q33DQ2.1 haplogroup. The most common second haplogroup in CBDA dominant cultivars is the 6 amino acid diverged form of Q33DQ2.1 (N90F, I266T, L370F, D420G, Y471C, T492S). Additionally, no CBDA dominant strains amplified with perfect Q8GTB6.1 active alleles with the MGC-2130 primers. The only two cultivars to amplify with the MGC-2130 primers had variants (Otto-A411V, and Blueberry Essence-P333R). The other three CBDA cultivars failed to amplify with MGC-2130 primers (WZ-CBD, Sour Tsunami, CannaTsu) despite gradient PCR optimization attempts.

This suggests UTR sequences of various THCAS haplogroups are associated with different chemotypes, and invokes an interest in the linked Bt:Bd allele described by de Meijer *et al*.). Weiblen *et al*. demonstrated this marker is tightly linked to THCAS and CBDAS. Pacifico *et al*. implied these B190/200 markers were imperfect in resolving the Bt:Bd from Bd:Bd alleles but never published the implied and improved B1080/1192 primers (Pacifico, 2006). Although, it is possible that the Bt allele is just in phase with the MGC-2130 primer set and predicts active form THCAS while the Bd allele is in phase with functional CBDAS, the current genome assemblies can only anchor partial B190 or B200 primer sequences (used to amplify the Bt:Bd allele) to a genomic contig (scaffold 19079). Sequencing these Bt:Bd products with Illumina sequencing demonstrated high coverage and high homology between the various sized bands suggesting a recently expanded repetitive element that is hard to assemble (Figure 7).

The UTR based MGC-2130 primers imply the THCAS replicated inactive events did not include 5 or 3’ UTRs. Primers that rest 387 & 160 bases back from the start and stop of the gene amplify fewer haplogroups of THCAS presumably due to the genomic divergence flanking the transcribed sequences. 10/13 of the cultivars amplified successfully with the more distant MGC-2130 primers (WZ-CBD, Sour Tsunami, and CannaTsu failed to amplify). Only one hybrid cultivar demonstrated a single haplogroup with both primers (Blueberry Essence Q8GTB6.1 P333R). The MGC-2130 primers appear to predominantly amplify the active forms and could possibly be used to better inform chemotype prediction. Due to the high polymorphism rate of this species and locus, sampling of additional cultivars with well-measured chemotypes is underway.

These haplogroups help to inform why copy number assays targeting highly conserved regions of THCAS can fail to differentiate active from inactive haplogroups. Cascini *et al*. demonstrate copy number variation in THCAS with qPCR but these primer selections would have failed to amplify only one inactive variant haplogroup in this study (Black84_Onofri-lbc1_C0_P0_NumRs92). A comprehensive sequencing strategy can properly identify active from inactive copies.

## Conclusions

The study of the inheritance of THCAS in modern drug type cultivars must consider polyploidy. Even though many of the replication events in THCAS appear to be pseudogenes or inactive forms, partially active forms are emerging and appear to be associated with more the balanced (Bt:Bd) THCA:CBDA cultivars. While de Meijer et al. demonstrate (Jurka & Kapitonov, 2001)chemotype prediction from the B190/B200 primers for the Bt:Bd allele, its precise location in the genome is difficult to uniquely resolve and while it can inform THCA and CBDA chemotype it has not been able to predict the magnitude of THCA or CBDA expression and some ambiguity on Bt:Bd vs Bd:Bd genotyping has been noticed. The sequencing of these alleles demonstrates a repetitive and polymorphic locus that appears difficult to design higher throughput qPCR assays for.

The observation of diverged THCAS genes and copy number variation may provide another tool in refining the chemotype prediction for *Cannabis* in light of previous work. Novel variants have been identified and phased with other variants of interest in THCAS. In light of both copy number variation, amino acid variation, B190-B200 alleles and van Bakels AAE3 observations, it is possible that the highly selected chemotype is experiencing convergent evolution. Of note are the several CWCTTA/TTAGWG “P Instability Factor” (PIF) sequences present in THCAS. While it is unclear if this replication event is transposon driven, the multiple, albeit small, recognition sequence signals in the gene and the repetitive sequence flanking THCAS is of note (Jurka & Kapitonov, 2001; Zhang *et al*., 2001). Structural variation in the human genome is often attributed to segmental duplication of ancestral alleles followed by Nonhomologous end joining (NHEJ) or non allelic homologous recombination (NAHR). VNTRs and retrotransposition are also known to drive these events (Kidd *et al*., 2008). While gene duplication of THCAS is common, the exact mechanism driving it remains to be determined.

## Methods

Two different amplicons for THCAS were selected. Primers described by Onofri *et al* were utilized but also redesigned as these previously described primers are internal to the gene (5’ ATG in the Forward primer is the start codon for the gene) and cannot inform on start codon or N-terminal signal peptide sequence. To multiplex multiple cultivars per SMRT cell, 5 prime dual 16 base pair molecular barcodes were utilized as described by the vendor.

### THCAS PCR

PCR was performed with 50ul 2x LongAmp (New England Biolabs- #M0287S), 8ul 10uM each primer, 42ul DNA+ddH20 (30ng total). Plant DNA was extracted with SenSATIVAx according to manufacturers instructions (Medicinal Genomics part #420001). DNA is eluted with 50ul ddH20.

THCAS was amplified from genomic 30ng of genomic DNA using primers Onofri-F (Forward;ATGAATTGCTCAGCATTTTCCTT) and Onofri-R (Reverse; ATGATGATGCGGTGGAAGA). Product was cycled using 98C 30 s followed by 28 cycles of 98C 10 s, 48C 30 s, 72C 90 s followed by a 72C for 5 min.

THCAS was also amplified from genomic 30ng of genomic DNA using primers MGC-2130-F (Forward; ATGGGAACCATAATAAACTATAAAAGTCATT) and MGC-2130-R (Reverse; TATGATTGTCTACACAGTTCTAATGTAGATTTTC). Product was cycled using 94C 30s followed by 28 cycles of 98C 10s, 58C 30s, 65C 150s followed by a 65C for 5 min.

PCR products were purified with equal volume of SenSATIVAx (100ul) followed by two 100ul 70% EtOH washes. Samples were eluted in 25ul ddH20 and quantified with Qubit and Agilent HS chips, normalized and pooled for Pacific Biosciences circular consensus sequencing. DNA barcodes recommended by the manufacturers were used to multiplex sequencing on each SMRT cell. Sequencing was performed by University of Florida service center.

## Nextera Libraries for Illumina phase confirmation

To generate sized shotgun libraries 9ul of THCAS PCR product (1.7ng/ul) was added to 20ul Nextera Buffer with 0.5ul Nextera Enzyme and 0.5ul ddH20. Nextera reaction was driven 7 minutes at 51C. Reaction was quickly purified with 40ul of SenSATIVAx. Tip mixes 10 times and placed on a magnet plate (Medicinal Genomics part#420202) for 10 minutes, removed supernatent and washed with 100ul 70% EtOH twice. After 5 minutes of benchtop room temp drying of residual EtOH the samples were eluted with 25ul ddH20.

## Post Nextera end healing or NT-PCR

To end repair the transposition event a nick translation event was performed prior to initial denaturization (NT-PCR). 31ul of LongAmp 2X master mix (New England Biolabs- #M0287S) was added to 6ul primer+indicies (2ul Index 1, 2ul Index 2, 2ul ILMN primers). 25ul Purified Nextera library DNA (62ul total reaction volume) was used for NT-PCR.

Reactions were cycled with 72C for 3 minutes, 98C for 30 seconds, 98C for 10 seconds, 63C for 30s, 72C for 1minute, Goto step 3 for 14cycles, 10C-hold. After NT-PCR, the sample was purified with 60ul SenSATIVAx added to the 62ul NT-PCR reaction. SenSATIVAx was tip mixed 10 times and placed on a magnet plate for 10 minutes. After separation and supernatant removal the sample was washed twice with 100ul of 70% EtOH. The beads were dried on a magnet plate for 10 minutes and eluted in 25ul ddH20

## SAGE size separation

25ul sample was added to 10ul R2 Marker (Sage Sciences) and loaded into a Sage Science Blue Pippin, 1.5% Agarose Dye-Free 40ul EM. Software was set to size cut at 600-800bp in size. 40-60ul of elution was captured in the output port. Optionally QC 1ul on an Agilent 2100 Bioanalyzer High Sensitivity chips. Loaded library according to Illumina V2 instructions protocol with 2x250bp reads.

## Bt:Bd Allele Amplification and sequencing

Primers described by de Meijer *et al*. were used named B190F(Forward; B190F TGCTCTGCCCAAAGTATCAA) and B200R (Reverse; CCACTCACCACTCCACCTTT).

Using 5ng input DNA was added to 25uL Q5 polymerase 2X Master Mix (NEB #). 1.25uL B190F Primer (10uM) with 1.25uL B200R Primer (10uM) were mixed with 22.5uL Water. Total volume was brought to 50ul with water + DNA (5ng total).

### Genomic amplification

PCR reactions were heated to 98C for 30 seconds and 40 cycles of 98C for 10s, 56C for 30s, 72C for 30s. A final 72C for 5 minutes was performed before 8C hold. PCR products were purified with 75ul SenSATIVAx. After two 100ul 70% EtOH washes, the beads were dried and eluted in 25ul ddH20. Yield was measured with a Qubit and an Agilent HS chip.

### PCR product Cloning

DNA libraries were constructed with 250ng DNA using NEB’s NEBNext Quick ligation module (NEB # E6056S). End Repair used 3ul of Enzyme Mix, 6.5ul of Reagent Mix, 55.5ul of DNA + ddH20. After End Repair, Ligation was performed directly with 15ul of Blunt End TA Mix, 2.5ul of Ilumina Adaptor (10uM) and 1ul of Ligation enhancer (assumed to be 20% PEG 6000). After 15-minute ligation at 25C, 3ul of USER enzyme was added to digest the hairpin adaptors and prepare for PCR. The USER enzyme was tipmixed and incubated at 37C for 20 minutes. After USER digestion, 86.5ul of SenSATIVAx was added and mixed. The samples were placed on a magnet for 15 minutes until the beads cleared and the supernatent could be removed. Beads were washed twice with 150ul of 70% EtOH. Beads were left for 10 minute to air dry and then eluted in 25ul of 10mM Tris-HCl.

### Amplification of libraries

Samples adapted with NEB adaptors were amplified with 25ul of 2X Q5 Polymerase (NEB). 1ul of 25uM i7 Primer and 1ul Universal primer were added. 23ul of Eluted DNA was added to make a 50ul total reaction. 98C for 30s was used to hot start the enzyme, followed by 6 cycles of 98C for 10s, 65C for 30s and 72C for 30s. A final 72C for 5 minutes was performed before 8C hold. PCR products were then purified using equal volumes (50ul) of SenSATIVAx. Samples were run on Qubit and Agilent to gauge their quality and further size selected on a SAGE Blue Pippin 1.2% Agarose dye free system to remove off target amplicons. (100-400bp). Quantified libraries were run on a MiSeq with V2 chemistry using 2x250bp reads.

### Analysis

Circular Consensus was achieved using Pacific Biosciences tools for de-multiplexing symmetric barcodes with the RS Long Amplicon Analysis.1 using SMRTanalysis 2.3.0 workflows (http://www.pacb.com/devnet/). Minimum size was selected as 1500 and 2300 for Onofri and MGC-2130 amplicons respectively with a maximum subread of 700. Fasta files were aligned with a CLCbio workstation and confirmed using BLAST to active and inactive accessions numbers (Q8GTB6.1 and Q33DQ2.1). Phylogenetic trees were graphed with CLC bio with E33090 set as the root.

Illumina reads were mapped with BWA-MEM version 0.7.12-r1044 to the E33090 with 600-650bp inserts selected. These reads were subsequently mapped to each PacBio derived haplogroup for each respective cultivar. Reads mapped to PacBio haplogroup references were displayed in the Integrated Genome Viewer or IGV (Robinson *et al*., 2011). Genbank accession numbers for this manuscript: SRP064442

## Author Contributions

KJM- Experimental design, barcoded PCR, Nextera, Sequencing and Manuscript drafting

YH- Barcoded PCR, Nextera, Sequencing, B190, B200 tests

VT- THCAS primer design and PacBio software installment.

SM- Illumina alignments to phased PacBio data and Figure generation

JS- Experimental optimizations and method development

LZ- Experimental optimizations and method development

DS- Manuscript review and drafting.

## Acknowledgments

Helpful communication from Ryan Lee, John McPartland, Brad Douglass, and Giuseppe Mandolino.

## References

Cascini F, Passerotti S, Boschi I. 2013. Analysis of THCA synthase gene expression in cannabis: a preliminary study by real-time quantitative PCR. Forensic Sci Int 231(1–3): 208–212.

Cascini F, Passerotti S, Martello S. 2012. A real-time PCR assay for the relative quantification of the tetrahydrocannabinolic acid (THCA) synthase gene in herbal Cannabis samples. Forensic Sci Int 217(1–3): 134–138.

Chaisson MJ, Huddleston J, Dennis MY, Sudmant PH, Malig M, Hormozdiari F, Antonacci F, Surti U, Sandstrom R, Boitano M, Landolin JM, Stamatoyannopoulos JA, Hunkapiller MW, Korlach J, Eichler EE. 2015. Resolving the complexity of the human genome using single-molecule sequencing. Nature 517(7536): 608–611.

de Meijer EP, Bagatta M, Carboni A, Crucitti P, Moliterni VM, Ranalli P, Mandolino G. 2003. The inheritance of chemical phenotype in Cannabis sativa L. Genetics 163(1): 335–346.

Eid J, Fehr A, Gray J, Luong K, Lyle J, Otto G, Peluso P, Rank D, Baybayan P, Bettman B, Bibillo A, Bjornson K, Chaudhuri B, Christians F, Cicero R, Clark S, Dalal R, Dewinter A, Dixon J, Foquet M, Gaertner A, Hardenbol P, Heiner C, Hester K, Holden D, Kearns G, Kong X, Kuse R, Lacroix Y, Lin S, Lundquist P, Ma C, Marks P, Maxham M, Murphy D, Park I, Pham T, Phillips M, Roy J, Sebra R, Shen G, Sorenson J, Tomaney A, Travers K, Trulson M, Vieceli J, Wegener J, Wu D, Yang A, Zaccarin D, Zhao P, Zhong F, Korlach J, Turner S. 2009. Real-time DNA sequencing from single polymerase molecules. Science 323(5910): 133–138.

Jurka J, Kapitonov VV. 2001. PIFs meet Tourists and Harbingers: a superfamily reunion. Proc Natl Acad Sci U S A 98(22): 12315–12316.

Kidd JM, Cooper GM, Donahue WF, Hayden HS, Sampas N, Graves T, Hansen N, Teague B, Alkan C, Antonacci F, Haugen E, Zerr T, Yamada NA, Tsang P, Newman TL, Tuzun E, Cheng Z, Ebling HM, Tusneem N, David R, Gillett W, Phelps KA, Weaver M, Saranga D, Brand A, Tao W, Gustafson E, McKernan K, Chen L, Malig M, Smith JD, Korn JM, McCarroll SA, Altshuler DA, Peiffer DA, Dorschner M, Stamatoyannopoulos J, Schwartz D, Nickerson DA, Mullikin JC, Wilson RK, Bruhn L, Olson MV, Kaul R, Smith DR, Eichler EE. 2008. Mapping and sequencing of structural variation from eight human genomes. Nature 453(7191): 56–64.

Kojoma M, Seki H, Yoshida S, Muranaka T. 2006. DNA polymorphisms in the tetrahydrocannabinolic acid (THCA) synthase gene in “drug-type” and “fiber-type” Cannabis sativa L. Forensic Sci Int 159(2–3): 132–140.

Mandolino C. 1999. Identification of DNA markers linked to the male sex in dioecious hemp (Cannabis sativa L.). Theor Appl Genet (1999) 98: 86Đ92.

McKernan KJ. https://archive.org/details/SequencingTheCannabisGenome) Sequencing The Cannabis Genome.

McKernan KJ. https://aws.amazon.com/datasets/the-cannabis-sativa-genome/. The Cannabis Sativa Genome.

Metzker ML. 2010. Sequencing technologies - the next generation. Nat Rev Genet 11(1): 31–46.

Nakamura K, Oshima T, Morimoto T, Ikeda S, Yoshikawa H, Shiwa Y, Ishikawa S, Linak MC, Hirai A, Takahashi H, Altaf-Ul-Amin M, Ogasawara N, Kanaya S. 2011. Sequence-specific error profile of Illumina sequencers. Nucleic Acids Res 39(13): e90.

Onofri C, de Meijer EP, Mandolino G. 2015. Sequence heterogeneity of cannabidiolic- and tetrahydrocannabinolic acid-synthase in Cannabis sativa L. and its relationship with chemical phenotype. Phytochemistry 116: 57–68.

Pacifico D. 2006. Genetics and marker-assisted selection of the chemotype in Cannabis sativa L. Molecular Breeding 17: 257–268.

Robinson JT, Thorvaldsdottir H, Winckler W, Guttman M, Lander ES, Getz G, Mesirov JP. 2011. Integrative genomics viewer. Nat Biotechnol 29(1): 24–26.

Shoyama Y, Tamada T, Kurihara K, Takeuchi A, Taura F, Arai S, Blaber M, Shoyama Y, Morimoto S, Kuroki R. 2012. Structure and function of 1-tetrahydrocannabinolic acid (THCA) synthase, the enzyme controlling the psychoactivity of Cannabis sativa. J Mol Biol 423(1): 96–105.

Sirikantaramas S, Morimoto S, Shoyama Y, Ishikawa Y, Wada Y, Shoyama Y, Taura F. 2004. The gene controlling marijuana psychoactivity: molecular cloning and heterologous expression of Delta1-tetrahydrocannabinolic acid synthase from Cannabis sativa L. J Biol Chem 279(38): 39767–39774.

van Bakel H, Stout JM, Cote AG, Tallon CM, Sharpe AG, Hughes TR, Page JE. 2011. The draft genome and transcriptome of Cannabis sativa. Genome Biol 12(10): R102.

Weiblen GD, Wenger JP, Craft KJ, ElSohly MA, Mehmedic Z, Treiber EL, Marks MD. 2015. Gene duplication and divergence affecting drug content in Cannabis sativa. New Phytol.

Zhang X, Feschotte C, Zhang Q, Jiang N, Eggleston WB, Wessler SR. 2001. P instability factor: an active maize transposon system associated with the amplification of Tourist-like MITEs and a new superfamily of transposases. Proc Natl Acad Sci USA 98(22): 12572–12577.

